# Age- and alcohol-related differences in adolescent neurometabolite levels

**DOI:** 10.1101/2025.07.25.666840

**Authors:** Maria I. Perica, Anna E. Kirkland, Louise Mewton, Lindsay M. Squeglia

## Abstract

Adolescence is a critical period for neurometabolite maturation as well as the onset of alcohol use, yet remains underexplored despite its significance for long-term neurodevelopmental outcomes. We used 3T proton magnetic resonance spectroscopy (MRS) to examine age- and alcohol-related associations with six neurometabolites in dorsal anterior cingulate cortex (dACC) that are involved in key neural functions: glutamate + glutamine (Glx), GABA plus macromolecules (GABA+), total N-acetylaspartate (tNAA), total choline (tCho), total creatine (tCr), and *myo*-inositol (mI). Participants were 84 adolescents (ages 17 – 22; 67% female) who completed MRS scans and self-reported past-60-day alcohol use via a modified Timeline Followback survey. Alcohol use variables included total drinking days, total binge drinking days, total number of drinks, and drinks per drinking day. Older adolescents had higher levels of GABA+, tNAA, tCho, and mI, and lower levels of Glx and Glx/GABA+; tCr was not associated with age. More alcohol use – specifically more drinking days, binge drinking days, and number of drinks – was associated with lower tNAA levels. Findings suggest age-related variation in dACC neurometabolites, which potentially reflect ongoing neuronal maturation, myelination, and shifts in excitatory and inhibitory neurotransmission. Lower tNAA among heavier drinkers may reflect associations between alcohol exposure and neuronal damage. Broader neurometabolic effects may emerge only with heavier or prolonged alcohol use.

## Introduction

Adolescence is a significant and distinct period of brain development, marked by dynamic neurobiological changes in regions critical for higher-order functions, such as cognition (1). Key neurodevelopmental processes during adolescence include thinning of cortical gray matter (2), pruning of synapses (3), myelination of axons (4), and changes in neurotransmitter systems (5,6). However, adolescence is also a time when heightened activity in neural reward systems can lead to increased risk-taking and sensation-seeking behaviors, such as alcohol use (7). In fact, adolescence is often when alcohol use is first initiated and begins to escalate notably (8). This presents a significant public health concern, as earlier initiation of alcohol use has been linked to more severe long-term consequences, including an increased risk of later meeting criteria for alcohol use disorder (AUD) (8–10). Moreover, alcohol use has been shown to significantly affect neurobiology, including neural processes that are actively developing during adolescence, which may lead to long-term, negative consequences for neurodevelopmental outcomes (11,12).

Despite these concerns, knowledge about how alcohol effects neurometabolite development in the human brain (e.g., neurotransmitter systems, neural function indicators) is limited. Much of the known research thus far has been conducted either in animal models, and has identified potential widespread neurometabolite alterations as a result of alcohol binging, withdrawal, and craving (13). Additionally, a recent meta-analysis of 43 magnetic resonance spectroscopy (MRS) studies investigating neurometabolite alterations as result of alcohol use found that the human literature also notes widespread neurometabolic impacts; however, the studies were disproportionately based on samples of middle-aged men, with no study samples under the average age of 21 (14). The overrepresentation of older, male samples limits our understanding of alcohol’s impact across sexes during adolescence – a key developmental period of potential neurometabolic development (6,15–19). Given the unique neurodevelopmental processes occurring during human adolescence, findings from adult samples or animal models may not generalize to the adolescent brain. Indeed, age has been identified as a crucial factor influencing the efficacy and safety of psychopharmacological treatments for psychiatric disorders (20), underscoring the need to account for developmental stage when considering the effects of alcohol on the brain and identifying neuroscience-informed treatment targets across the lifespan.

Here, we aimed to address several gaps in the literature. First, we explored how a range of neurometabolites – each supporting critical neural functions – develop during adolescence. While the relationship between neurometabolite levels and age has been increasingly studied across the lifespan, particularly early development and older adulthood, adolescence remains understudied despite the significance of this developmental stage (21,22). Second, we characterized how early alcohol use in adolescence alters neurometabolite levels to understand whether these effects align with or diverge from the predominantly adult literature. We applied 3T proton magnetic resonance spectroscopy (¹H-MRS) in 84 participants aged 17 – 22 to investigate 1) age-related differences in neurometabolites in the dorsal anterior cingulate cortex (dACC) and 2) associations between alcohol use and dACC neurometabolite levels. We selected dACC as our region of interest as it continues to develop late into adolescence, it plays a role in higher-order cognitive functions, and it has shown to be impacted by youth binge drinking (23–25). Additionally, we chose to assess all metabolites commonly quantified from ^1^H-MRS data, each of which has been linked to key neural functions: N-acetylaspartate (tNAA; marker for neuronal health, integrity, density), choline (tCho; putative marker for lipid membrane synthesis/turnover), GABA (GABA+; inhibitory neurotransmitter), myo-inositol (mI; putative marker for lipid membranes and glia), glutamate + glutamine (Glx; excitatory neurotransmitter glutamate + glutamine), and creatine (tCr; marker for energetic/metabolic activity). We additionally investigated the ratio between Glx and GABA+, as this ratio may be related to the excitatory/inhibitory balance in the brain, which is thought to be shifting toward inhibition through adolescence (5). Findings from this study will enhance our understanding of how these neurometabolites develop through adolescence, as well as how alcohol use may affect them.

## Methods and Materials

### Participants

This is a secondary data analysis of two completed double-blind, placebo-controlled, crossover clinical trials assessing the neural effects of cannabidiol (NCT05317546) or N-acetylcysteine (NCT03238300) in adolescents who use alcohol (26, 27). The total sample across both studies was 84 participants from 17 – 22 years old (mean age = 19.5 years old, 65% female). Inclusion and exclusion criteria are briefly described here; for full inclusion and exclusion criteria for both studies see Supplementary Table 1 and see Supplementary Table 2 for sample demographics for each study. For the cannabidiol study, participants (n= 34; 17 – 22 years old) all met criteria for AUD within the past year, had at least 1 AUD symptom in the past 30 days (excluding craving), and had consumed alcohol in the past 2 weeks prior to screening. Participants in the cannabidiol study completed one scan session with placebo and one with cannabidiol, and only data from the placebo condition was used for this analysis. No carry-over effects were found between scanning sessions in this study, suggestive of a sufficiently long washout period to clear cannabidiol before receiving placebo (27). For the N-acetylcysteine study, participants included adolescents (n = 50; 17 – 19 years old) who reported alcohol use, including heavy alcohol use, but did not necessarily meet criteria for AUD (n = 30 with AUD; n = 20 without AUD) (26). Heavy alcohol use was defined as at least 4 drinking occasions with at least 3 drinks per occasion over 90 days prior to participating in the study (28). Participants in the N-acetylcysteine study completed a baseline scan prior to being randomized to N-acetylcysteine or placebo, and only baseline data was used for this analysis.

Exclusion criteria for both studies included: 1) history of a significant or acutely unstable medical, neurological, psychiatric, or substance use problems (other than alcohol use) as assessed with the Mini-International Neuropsychiatric Interview; 2) history of neurodevelopmental condition that could impact brain development; 3) currently pregnant, trying to become pregnant, or breastfeeding; 4) positive urine toxicology screen for narcotics, amphetamines, sedatives, hypnotics, or opiates at screening; 5) history of treatment or treatment-seeking for alcohol use; 6) a score of 10 or higher on the Clinical Institute Withdrawal Assessment for Alcohol; 7) acute drunkenness or consumption of alcohol within 12 hours of visit; and 8) MRI contraindications (e.g., braces, claustrophobia, irremovable metal implants or piercings). For participants under the age of 18, parental consent and participant assent were obtained prior to collecting data. All participants were recruited from the community using mixed-methods, and they were financially compensated for their participation. All experimental procedures were approved by the Medical University of South Carolina Institutional Review Board.

### Proton Magnetic Resonance Spectroscopy (1H-MRS) Data Acquisition

Both studies used the same 1H-MRS protocols. All scans were performed on the same Siemens 3.0T Prismafit MR scanner with an actively shielded magnet and high-performance gradients (80 mT/m, 200 T/m-sec) using a 32-channel head coil. High-resolution structural scans were acquired using a magnetization prepared rapid gradient echo (MPRAGE) sequence to allow for later registration to a predefined region-of-interest (ROI), the dorsal anterior cingulate cortex (dACC) (scan parameters: TR/TE = 2250/4.18 ms; flip angle = 9°; field of view = 256 mm^2^; voxel size= 1 mm^2^; 176 contiguous 1-mm-thick slices). The MRS protocol used was based on previously published studies (29,30). The dACC voxel was placed on midsagittal T1-weighted images (Figure 1), anterior to the genu of the corpus callosum, with the ventral edge of the voxel aligned with the dorsal edge of the genu (31), with a voxel size of (30 x 25 x 25) mm^3^ (31,32). Following FAST(EST)MAP shimming (33), single-voxel water-suppressed 1H-MRS spectra were acquired (water suppression bandwidth: 50 Hz for Glx and GSH or 100 Hz for GABA+, spectral bandwidth 2000 Hz, 1024 spectral points) 1H-MRS spectra were acquired with the following sequences: 1) SIEMENS Point Resolved Spectroscopy (PRESS) sequence: TR=2000 ms; TE=40 ms ; number of averages = 256; 2) SIEMENS WIP MEGA-PRESS sequence for GABA+: Edit ON(OFF)=1.90 (7.46) ppm; TR=2000 ms, TE=68 ms; number of averages=160 (Supplementary Figure 1). Unsuppressed water spectra were co-acquired and scaled for partial volume effects and relaxation and used as a concentration reference. Six saturation bands (41-mm thickness) were placed for outer volume suppression at a distance of 0.8 cm from each voxel face.

**Figure 1.**
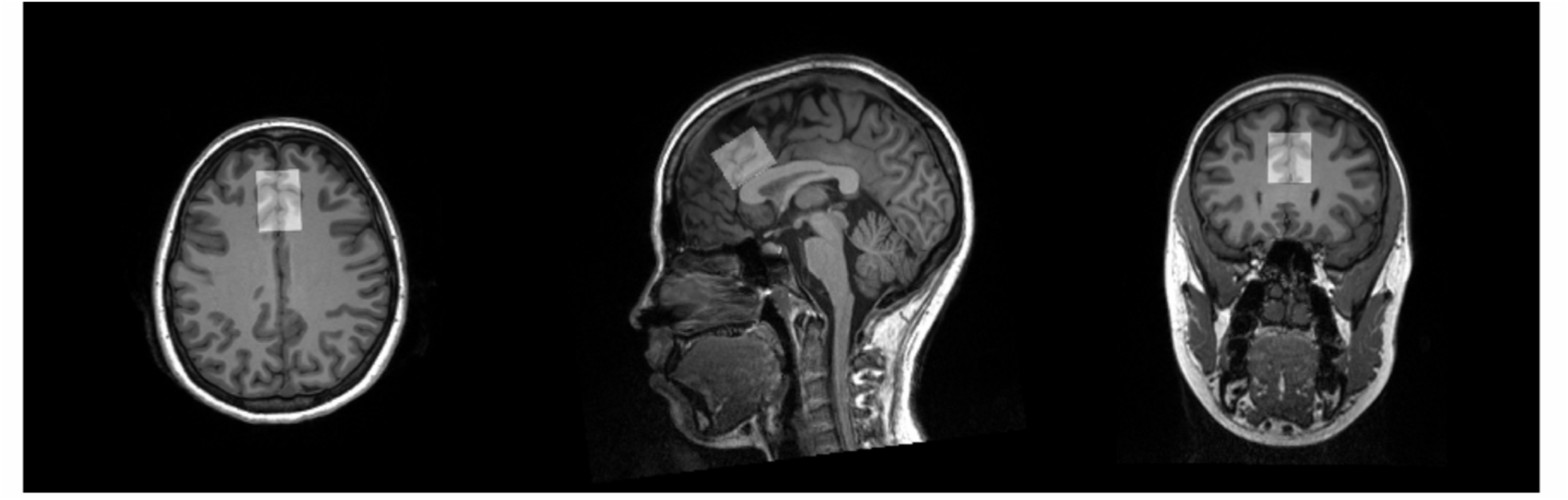
Example of typical voxel placement in dACC on a T1 image.

### Quantification of Metabolite Levels and Data Quality Criteria

Osprey software was used to quantify levels of metabolites from 1H-MRS data (34). Standard parameters were used for PRESS processing (metabolite fit range: 0.2 to 4.2 ppm; water fit range: 2.0 to 7.4 ppm; knot spacing=0.4), and parameters were optimized for GABA+ processing (metabolite fit range: 0.5 to 4.0 ppm; water fit range: 2.0 to 7.4 ppm; knot spacing=0.55; MM09 hard modelling) (35). Segmentation of T1-weighted images was done using SPM12 (36) to quantify voxel tissue composition (fraction of gray matter [GM], white matter [WM], and cerebrospinal fluid). All metabolite levels are expressed in fully tissue-and-relaxation-corrected molal concentrations (mol/kg) (37).

Data signal-to-noise ratio (ratio between amplitude of Cr peak and standard deviation of detrended noise), linewidth for Cr (full-width half-maximum [FWHM] of single-Lorentzian fit to Cr peak), and linewidth for water (H2O; FWHM of single-Lorentzian fit to H2O reference peak) was used as quality criteria. Individual cases were excluded from analyses if the Cr SNR was >3 standard deviations from the group mean or if the linewidth was >11 Hz.

### Alcohol Use Data

Alcohol and other substance use (e.g.., cannabis use, nicotine use) data were self-reported by participants using the modified Timeline Follow-Back (TLFB) survey (38) with a reporting window over the past 60 days from their screening visit. Of note, while the N-acetylcysteine study collected 90-days of TLFB data at screening, we only report days 1-60 here to match the cannabidiol study procedures. The following substance use variables were calculated and used in analyses as variables of interest and covariates: 1) total number of drinking days, 2) total number of standard drinks consumed, 3) total number of binge drinking days (4+ drinks for females, 5+ drinks for males per day), 4) maximum number of drinks consumed on a single drinking day, 5) average number of drinks per drinking day (DPDD), 6) nicotine use days, and 7) cannabis use days (summed over various methods of use for nicotine and cannabis; see Supplementary Figure 2 for correlations between variables).

### Statistical Analysis

All analyses were conducted in R Statistical Software (39). To investigate associations between metabolites and age (Aim 1), we used linear regression. To test for nonlinear age-related associations, we used Generalized Additive Models (GAM) and Akaike’s Information Criterion (AIC) to compare the model with the nonlinear (smooth) age term to the model with the linear age term (40,41). In all models, a covariate was included to control for individual differences in tissue composition of the dACC voxel, the ratio of gray matter (GM) to brain matter (BM) in the voxel (GM/BM). To investigate the effect of alcohol use on metabolites (Aim 2), we used linear regression with covariates for GM/BM and age. Inclusion of other covariates was tested using the likelihood-ratio test (42). Covariates tested included sex, number of cannabis use days and number of nicotine use days, to ensure that any findings were specific to alcohol use rather than general substance use. Covariates were included in the model if they made a significant difference to the model. Due to the exploratory nature of this study, p-values reported were not corrected for multiple comparisons (43).

## Results

### Participants

84 participants were included ranging from 17 – 22 years of age (mean age = 19.54, SD = 1.32). 67% of the recruited sample reported their sex as female and 90% of the sample reported their race as white. 76% of the overall sample met criteria for AUD in the past year, with 45% meeting criteria for mild AUD, 19% meeting criteria for moderate AUD, and 12% meeting criteria for severe AUD. See Table 1 for complete sample characteristics.

**Table 1.** Sample characteristics.

|  | N= 84 total |
| --- | --- |
| **Age**, range<br>mean (SD) | 17 – 22<br>19.54 (1.32) |
| **Sex**, n (%)<sup>1</sup><br>Female<br>Male | 56 (67%)<br>28 (33%) |
| **Race**, n (%)<br>Asian<br>Black/African American<br>White | 3 (4%)<br>5 (6%)<br>76 (90%) |
| **Ethnicity**, n (%)<br>Hispanic/Latino<br>Not Hispanic/Latino | 2 (2%)<br>82 (98%) |
| **Grade**, n (%)<br>10<sup>th</sup> grade<br>11<sup>th</sup> grade<br>12<sup>th</sup> grade/GED equivalent<br>Some college<br>Bachelor's Degree | 2 (2%)<br>8 (10%)<br>33 (40%)<br>34 (40%)<br>7 (8%) |
| **Mental Health Disorders**<sup>2</sup>, n (%)<br>Attention Deficit Hyperactivity Disorder<br>Obsessive-Compulsive Disorder<br>Major Depressive Episode<br>Generalized Anxiety Disorder<br>Social Anxiety Disorder<br>Panic Disorder<br>Alcohol Use Disorder<br>Mild<br>Moderate<br>Severe<br>Cannabis Use Disorder<br>Mild<br>Moderate<br>Severe | 13 (15%)<br>2 (2%)<br>4 (4%)<br>8 (9%)<br>3 (3%)<br>4 (4%)<br>64 (76%)<br>38 (45%)<br>16 (19%)<br>10 (12%)<br>32 (38%)<br>14 (17%)<br>12 (14%)<br>6 (7%) |
| **Alcohol Use**<sup>3</sup><br>Participants with alcohol use in past 60 days n (%)<br>Number of drinking days, mean (SD)<br>Total number of standard drinks, mean (SD)<br>Number of binge drinking days, mean (SD)<br>Maximum number of drinks in one day, mean (SD)<br>Number of standard drinks per drinking day, mean (SD)<br>Age of first full drink<br>Past 24 hrs self-reported alcohol use<br>Past 12 hrs self-reported alcohol use | 84 (100%)<br>14.49 (6.71)<br>68.49 (46.58)<br>8.29 (6.41)<br>9.38 (5.50)<br>4.67 (2.04)<br>15.55 (1.67)<br>13 (15%)<br>0 (0%) |
| **Other Substance Use**<sup>3</sup><br>Participants with cannabis use in past 60 days<br>Number of cannabis use days, mean (SD)<br>Urine tox screen positive for THC<br>Participants with nicotine<sup>4</sup> use in past 60 days<br>Number of nicotine use days, mean (SD) | 54 (64%)<br>10.32 (17.11)<br>17 (20%)<br>48 (57%)<br>17.95 (23.30) |

Note: <sup>1</sup>Percents are rounded to the nearest whole number. <sup>2</sup>MINI = Mini International Neuropsychiatric Interview; Current diagnosis was used ensuring that all diagnoses reflected participants conditions at the time of assessment. Only diagnoses assessed between both samples were reported.<sup>3</sup> Based on 60-day Timeline Follow Back (TLFB). Number of substance use days are only quantified using participants that use that substance. <sup>4</sup>Any nicotine use, including tobacco and e-cigarettes.

### Aim 1: Age-related associations with metabolite levels

We characterized age-related differences in tNAA, tCho, GABA+, mI, Glx, and tCr, covarying for GM/BM in the dACC voxel. We additionally created a variable of the ratio between Glx and GABA+ to directly investigate age-related differences in the Glx/GABA+ balance or excitatory/inhibitory balance. For all models, GAM models did not outperform linear regression models by more than 2 AIC units (Supplementary Table 3-4), so linear regression models were used for all models going forward for simplicity and interpretability.

Results showed that older adolescents had higher levels of tNAA (Standardized coefficient *β* = 0.35, *p*=0.001), tCho (*β* = 0.35, *p*<0.001), GABA+ (*β* = 0.45, *p*<0.001) and mI (*β* = 0.38, *p*<0.001) (Figure 2; Table 2). Results showed that older adolescents also had lower levels of Glx (*β* = -0.40, *p*<0.001) as well as a lower Glx/GABA+ ratio (*β* = -0.48, *p*<0.001). Finally, results also showed no association with age for tCr levels (*β* = -0.17, *p*=0.13).

**Figure 2.**
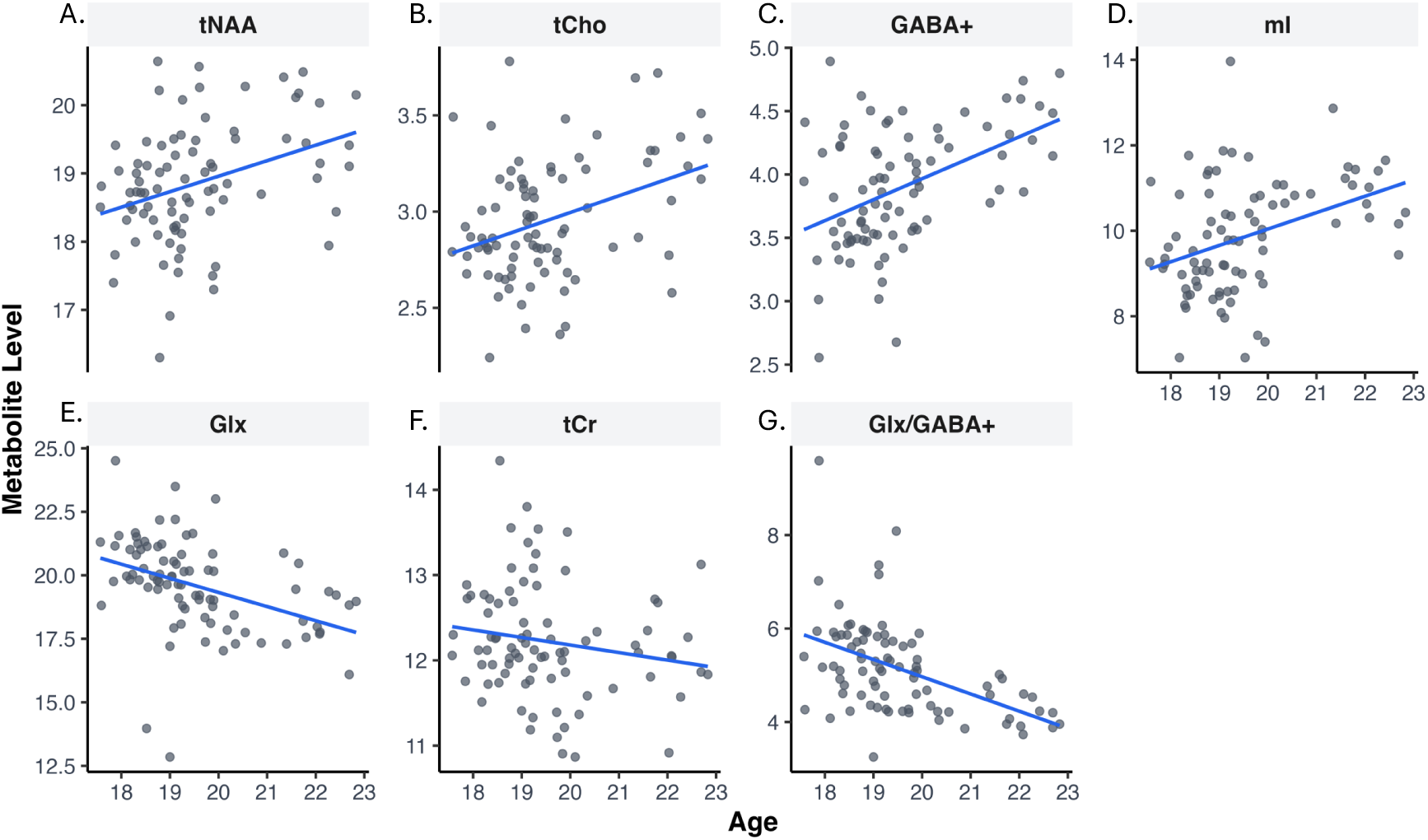
Association between metabolites and age with linear model fits. Scatter plots shown correlation between age and A) total N-acetylaspartate-containing compounds (tNAA). B) total choline-containing compounds (tCho). C) GABA + macromolecules (GABA+). D) *myo-*Inositol (mI). E) Glutamate + glutamine (Glx). F) Total creatine-containing compounds (tCr). G) Glx/GABA+ ratio (Glx/GABA+).

**Table 2.** Linear model output for associations between neurometabolites and age.

| Metabolite | Predictor | $\beta$ | $\beta$<br>95% CI<br>[LL, UL] | t | p |
| --- | --- | --- | --- | --- | --- |
| tNAA | **Age** | 0.35 | [0.15, 0.56] | 3.39 | 0.0011** |
|  | GM/BM | -0.11 | [-0.32, 0.09] | -1.09 | 0.28 |
| tCho | **Age** | 0.35 | [0.15, 0.56] | 3.44 | <.001** |
|  | GM/BM | 0.15 | [-0.05, 0.36] | 1.48 | 0.14 |
| GABA+ | **Age** | 0.45 | [0.25, 0.65] | 4.56 | <0.001** |
|  | GM/BM | 0.08 | [-0.11, 0.28] | 0.85 | 0.40 |
| ml | **Age** | 0.38 | [0.18, 0.58] | 3.70 | <0.001** |
|  | GM/BM | 0.07 | [-0.13, 0.28] | 0.73 | 0.47 |
| Glx | **Age** | -0.40 | [-0.60, -0.20] | -3.94 | <0.001** |
|  | GM/BM | 0.00 | [-0.20, 0.21] | 0.034 | 0.97 |
| tCr | Age | -0.17 | [-0.39, 0.05] | -1.55 | 0.13 |
|  | GM/BM | -0.10 | [-0.31, 0.12] | -0.88 | 0.38 |
| Glx/GABA+ | Age | -0.48 | [-0.67, -0.29] | -4.92 | <0.001** |
|  | GM/BM | -0.05 | [-0.24, 0.15] | -0.49 | 0.62 |

Note: $\beta$ indicates standardized coefficients. LL and UL indicate the lower and upper limits of a confidence interval, respectively. Bolded predictor variables represent $p < 0.05$ . \* indicates $p < 0.05$ . \*\* indicates $p < 0.01$ .

### Aim 2: Alcohol-related associations with metabolite levels

We then assessed how variation in alcohol use was associated with metabolite levels. For all base models, we included GM/BM and age as covariates. We additionally used the likelihood ratio test to test whether including sex, number of cannabis use days, or number of nicotine use days would improve model fit. For all models, we found that these covariates did not improve fit above the base model, and thus these covariates were not included going forward (Supplementary Table 5). Additionally, we found no associations between age and all our alcohol variables (*p* values > 0.05).

Results showed that levels of tNAA were associated with self-reported alcohol use (Table 3). Specifically, there were lower levels of tNAA with higher numbers of drinking days (*β* = -0.26, *p*=0.012), higher numbers of total drinks consumed (*β* = -0.21, *p*=0.043), and higher numbers of binge drinking days (*β* = -0.22, *p*=0.037) (Figure 3). No other metabolites were associated with alcohol use (Supplementary Table 6).

**Figure 3.**
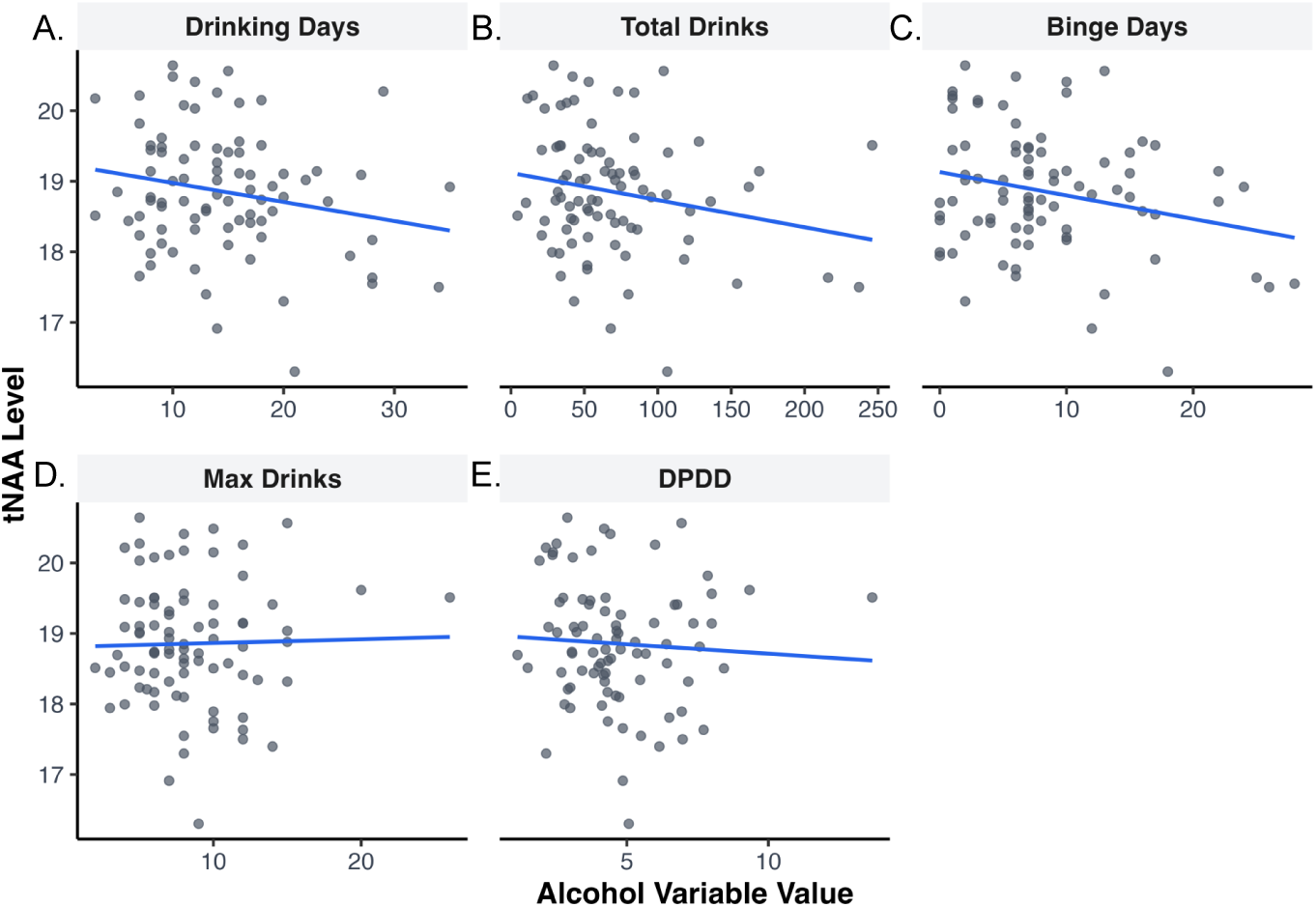
Association between alcohol (past 60 days) and total N-acetylaspartate-containing compounds (tNAA) with linear model fits. A) Days spent drinking alcohol. B) Total drinks consumed. C) Maximum number of drinks consumed in one drinking episode. D) Drinks per drinking day. E) Binge drinking episodes.

**Table 3.**
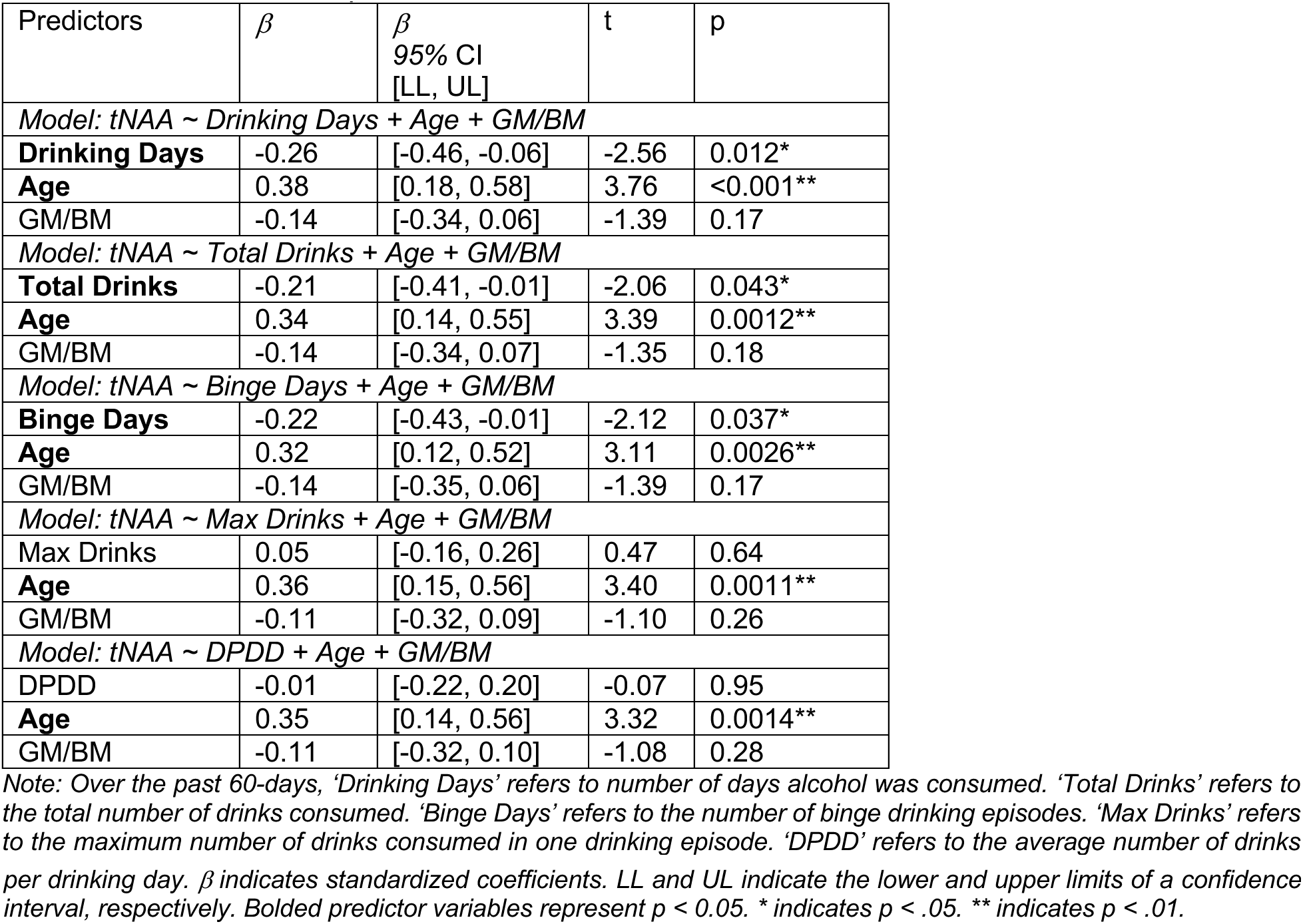
Linear model output for associations between tNAA and alcohol variables.

| Predictors | $\beta$ | $\beta$<br>95% CI<br>[LL, UL] | t | p |
| --- | --- | --- | --- | --- |
| *Model: tNAA ~ Drinking Days + Age + GM/BM* |  |  |  |  |
| **Drinking Days** | -0.26 | [-0.46, -0.06] | -2.56 | 0.012* |
| **Age** | 0.38 | [0.18, 0.58] | 3.76 | <0.001** |
| GM/BM | -0.14 | [-0.34, 0.06] | -1.39 | 0.17 |
| *Model: tNAA ~ Total Drinks + Age + GM/BM* |  |  |  |  |
| **Total Drinks** | -0.21 | [-0.41, -0.01] | -2.06 | 0.043* |
| **Age** | 0.34 | [0.14, 0.55] | 3.39 | 0.0012** |
| GM/BM | -0.14 | [-0.34, 0.07] | -1.35 | 0.18 |
| *Model: tNAA ~ Binge Days + Age + GM/BM* |  |  |  |  |
| **Binge Days** | -0.22 | [-0.43, -0.01] | -2.12 | 0.037* |
| **Age** | 0.32 | [0.12, 0.52] | 3.11 | 0.0026** |
| GM/BM | -0.14 | [-0.35, 0.06] | -1.39 | 0.17 |
| *Model: tNAA ~ Max Drinks + Age + GM/BM* |  |  |  |  |
| Max Drinks | 0.05 | [-0.16, 0.26] | 0.47 | 0.64 |
| **Age** | 0.36 | [0.15, 0.56] | 3.40 | 0.0011** |
| GM/BM | -0.11 | [-0.32, 0.09] | -1.10 | 0.26 |
| *Model: tNAA ~ DPDD + Age + GM/BM* |  |  |  |  |
| DPDD | -0.01 | [-0.22, 0.20] | -0.07 | 0.95 |
| **Age** | 0.35 | [0.14, 0.56] | 3.32 | 0.0014** |
| GM/BM | -0.11 | [-0.32, 0.10] | -1.08 | 0.28 |

Note: Over the past 60-days, 'Drinking Days' refers to number of days alcohol was consumed. 'Total Drinks' refers to the total number of drinks consumed. 'Binge Days' refers to the number of binge drinking episodes. 'Max Drinks' refers to the maximum number of drinks consumed in one drinking episode. 'DPDD' refers to the average number of drinks
per drinking day. $\beta$ indicates standardized coefficients. LL and UL indicate the lower and upper limits of a confidence interval, respectively. Bolded predictor variables represent $p < 0.05$ . \* indicates $p < .05$ . \*\* indicates $p < .01$ .

## Discussion

In this study, we provide novel evidence of age-related differences in neurometabolites within the dACC during late adolescence, as well as the associations between alcohol use and neurometabolite levels. We identified that older adolescents had higher levels of tNAA, tCho, GABA+, and mI, alongside lower levels of Glx. We also find that the Glx/GABA+ ratio is lower in older adolescents. In contrast, tCr levels did not vary with age. Notably, we additionally found that more alcohol use was associated with lower tNAA levels, in direct opposition to the positive association between age and tNAA that we observed. Interestingly, we did not find any significant associations between alcohol use and the other metabolites we investigated. Further, cannabis and nicotine use did not show any significant effects on neurometabolites over and above the effects of alcohol use. These findings offer new insights into the age-related variation in dACC neurometabolite levels during adolescence, and highlight tNAA as a potential early marker of neuronal damage linked to alcohol use.

Our results contribute evidence toward the understanding of neurometabolite levels during adolescence in the dACC, while also replicating and extending prior work. Our findings of lower levels of Glx, higher levels of GABA+, and a lower ratio of Glx/GABA+ with older age through adolescence aligns with findings from preclinical animal model literature suggestive of a shift in the balance between excitatory and inhibitory neurotransmission during adolescence in executive brain regions, such as dACC, which could support cognitive and socioemotional development (5). Additionally, some more recent MRS studies in human adolescents have also found lower dACC glutamate levels in older adolescents/adults relative to younger adolescents (6,17,19). In the context of adolescence, decreases in MRS glutamate-related metabolites can potentially be understood as resulting from known developmental mechanisms such as pruning of primarily excitatory synapses (44,3,6). However, other studies have not observed age-related associations in dACC glutamate (15,16,18). We also found higher levels of dACC GABA+ in older adolescents, consistent with a study reporting higher GABA/Cr levels in older adolescents/young adults (18 – 24 years old) compared to earlier in adolescence (12 – 14 years old) (18). However, it should be noted that other studies have reported null findings for age-related association with GABA/Cr or GABA+ in dACC (6,19). Finally, developmental studies that have looked at the ratio of glutamate to GABA in frontal brain regions are sparse, but one 7T study found no associations between age and Glu/GABA ratio in dACC (6), in contrast with our findings of a lower Glx/GABA+ ratio in older adolescents. This could be due to a variety of methodological differences from sample size and composition (non-alcohol using youth in Perica et al., 2024 vs alcohol-using in the present manuscript) to acquisition to processing pipeline. This underscores the need for further replication, as well as larger, longitudinal studies to clarify these within-person trajectories. These large-sample studies are also needed to clarify baseline trajectories of Glx, GABA, and other metabolites in non-alcohol-using youth in order to better distinguish normative developmental trajectories from alcohol-related effects.

Far less is known about the developmental trajectories of other neurometabolites in the dACC during adolescence. In our sample, we found a positive association between age and tNAA levels. NAA is abundant in neurons, but despite that, its specific function remains obscure. NAA has been thought to be a marker of neuronal, axonal, and dendritic viability, as reduced NAA levels are consistently observed in neurodegenerative and neuropsychiatric disorders where neuronal loss or dysfunction is present (45). Therefore, it is possible that age-related increases in tNAA during adolescence may reflect the development of synapses, dendrites, and neurons (46). Consistent with our findings in 17 – 22 year olds, another study found age-related increases in dACC NAA in a sample of 234 9 – 12 year olds (47). However, two other studies have reported group comparisons in dACC tNAA during the adolescent period, and neither found differences between younger adolescent and older adolescent/adult age groups (17,18). These studies had smaller adolescent samples (n = 11 and n = 30 respectively) and may have been underpowered to detect effects. Additionally, these studies normalized their tNAA metabolite values to creatine, which our study did not, in addition to using categorical age group comparisons rather than modeling age continuously, potentially limiting the ability to detect more subtle developmental variation. Our findings add to this limited literature by providing evidence for ongoing maturation of tNAA during adolescence. Future work should aim to further uncover the role of tNAA in adolescent neurodevelopmental processes.

Similarly, relatively little research has investigated tCho or mI in dACC during adolescence. Here, we find that both metabolites have higher levels in older adolescents compared to younger adolescents. Both Cho and mI are thought to be markers of processes related to lipid membranes, in addition to mI being thought of as a marker for glial cells (48). Increases during the adolescent period may reflect changes in lipid membranes, such as known developmental increases in myelination of axons during adolescence, or maturation of glia (4,48). However, as with NAA, the function of both metabolites remains to be more clearly understood, especially their roles in neurodevelopment. To our knowledge, only two studies have reported on tCho through adolescence with mixed results, where one study found higher levels of tCho in a group of 9 – 12 year olds with age (47), and another study reported no difference between two groups of younger and older adolescents in tCho/Cr or mI/ Cr (18).

Finally, we find no age-related associations in tCr through adolescence. To our knowledge, only one other study has reported age effects for tCr in dACC, and this study found higher tCr levels with age in a group of 9 – 12 year olds (47). The limited literature on tCr likely stems from its common use as a reference metabolite for normalization, rather than as a primary focus of study. However, tCr plays a critical role in energetic and metabolic processes in the brain and more research is needed to understand the role it may be playing during development (48). Our use of water-referencing in this study allows us to study tCr directly and contributes novel information about tCr during later adolescence. Further, referencing to Cr as a general practice presents a number of issues, such as increasing the variability and noise when creating ratios of metabolites to tCr (48–50). Therefore, our use of water-referenced values contributes valuable information to the literature about all our reported neurometabolites.

Our findings also provide novel information about the effect of alcohol use on neurometabolite levels during adolescence, specifically that more drinking days, binge drinking, and total number of drinks consumed is associated with lower NAA levels. The literature thus far has skewed heavily toward studying the impact of alcohol use on the brain in middle-aged, mostly male samples (14). A prior study with a portion of the dataset used in this secondary data analysis did not find any differences in any neurometabolite levels between an alcohol-using group and non-alcohol-using group (51). However, this study only included adolescents ages 17-19, which may be too young to see effects. Further, by combining two datasets in this study, one which included participants diagnosed with AUD, we may have been better powered to detect the effects seen with tNAA due to a larger sample as well as greater range in severity of alcohol use. Indeed, our findings are in line with another study of binge drinking in 18 – 24 year olds that found that greater binge drinking was associated with lower NAA levels in dACC (52). Our findings of lower NAA levels in dACC as a result of alcohol use is also consistent with the broader adult literature(14). However, in contrast to our findings, two studies did find effects of adolescent binge drinking on GABA levels, where more binge drinking was associated with lower GABA levels (52,53). It should be noted that this could be due to methodological differences, as GABA is less reliably measured than NAA due to the more dynamic nature of GABA synthesis as well as its low concentration in the brain (54). Further, in contrast with our study, the average age in those studies with slightly older (22.3 years old and 21.9 years old respectively), suggesting that more widespread neurometabolic effects may only emerge later in development.

This interpretation aligns with findings from the previously mentioned meta-analysis of primarily adult studies, which found widespread effects of alcohol use on neurometabolites (such as lower GABA and NAA levels in the dACC), suggesting that broader neurometabolic impacts may occur only after more prolonged alcohol use or later in the lifespan. It is possible that the highly plastic nature of the younger adolescent brain may allow for greater recovery potential (5), thus resulting in fewer long-lasting neural changes following substance and alcohol use. However, given the remarkable consistency of lower NAA levels as a result of alcohol use in our study, as well as the one other adolescent alcohol use study and the broader adult literature, NAA may have value as a potential early biomarker or treatment target warranting further investigation. Lower levels of NAA may represent neuronal damage as a result of alcohol use, as lower NAA levels are often seen in a variety of neurodegenerative disorders (45). Further, it is known that alcohol has neurodegenerative effects in youth, including decreased gray matter volume and cortical thickness (55). However, given the cross-sectional nature of this study and the majority of the current literature, it is difficult to interpret lower NAA levels associated with alcohol use. These lower levels may represent neuronal damage resulting from alcohol use or they could represent maturational delay, as we find higher levels of NAA in older adolescents, or they could be a result of a third variable correlated with alcohol use. Longitudinal studies are needed to disentangle the temporal nature of these remaining questions. Further, the exact function of NAA in dACC and what lower levels vs higher levels of NAA represent in the brain broadly remains to be fully elucidated. Uncovering more about the role of NAA during neurodevelopment may help inform targeted treatment research. Finally, in line with our findings here, the extant primarily adult literature does not find impacts on Glx, tCho, tCr, or mI (14). More studies are needed to replicate null findings or to investigate these metabolites in the first place, especially during development.

Notably, although we systematically tested both cannabis use days and nicotine use days, neither emerged as significantly effecting neurometabolite levels beyond the impact of alcohol use. A systematic review of alcohol and cannabis use during adolescence on structural and functional neurodevelopment found more pronounced neural effects relating to alcohol use at lower levels than to cannabis use (55). However, neural effects of cannabis use and nicotine use have certainly been noted (56–58). Our lack of findings here may have been due to being underpowered to detect them in this study as not all of our participants reported cannabis or nicotine use.

## Strengths and Limitations

This study has several notable strengths. First, this study’s inclusion of multiple neurometabolites provides a more comprehensive view of neurochemical development during adolescence. Our use of water-referencing in our MRS data to quantify neurometabolites overcomes some of the limitations caused by normalizing to creatine, as is commonly done across the MRS literature. Further, a key strength of this study is the use of a relatively large and developmentally younger sample compared to many previous MRS studies both of development and of alcohol use, enhancing the generalizability of our findings to alcohol exposure in youth broadly. Importantly, our use of an adolescent sample allowed us to highlight NAA as a specific and early marker of alcohol’s effects on the brain, which differs from more widespread findings in the adult literature and highlights the need for further research into NAA and its role in neurodevelopment.

However, several limitations should be noted. First, we present cross-sectional data, which limits our ability to draw causal inferences or assess within-person change. Second, we only present data from one brain region which, although common in MRS studies that have to use relatively large voxel sizes, limits our ability to understand impacts on other regions or broader networks. Longitudinal studies that include a variety of brain regions are needed to determine whether alcohol use leads to prospective alterations in neurometabolite trajectories over the course of development. Third, although alcohol use is normative and widespread among adolescents in the United States (59), our sample included only adolescents that reported some amount of alcohol us. Future studies should aim to build on our findings with longitudinal data from non-alcohol-using samples to help further clarify baseline developmental trajectories of all of these critical neurometabolites. Fourth, our overall sample is predominantly female (67%), which presents some concerns to both generalizability, as well as potential confounds, such as fluctuating hormonal differences that can impact measured metabolite levels (60). However, substance use literature has historically overrepresented white men, and thus predominantly female samples provide valuable novel information as well. Finally, although our sample size is larger than in many prior studies, replication in larger and more diverse samples will be needed to confirm and extend these findings to the broader population.

## Conclusions

Our study provides valuable information about neurometabolite variation in adolescence, in addition to associations between alcohol use and neurometabolite levels during adolescence. We report a variety of age-related differences in several neurometabolites, with older adolescents having higher levels of tNAA, tCho, GABA+, and mI, and lower levels of Glx, while tCr levels did not vary with age. In addition, we find that the Glx/GABA+ ratio is lower with older age. Further, alcohol use during adolescence was associated with lower levels of tNAA specifically, but not other metabolites. The contrast between the age-related increase in tNAA and alcohol-related decrease in tNAA suggests that alcohol use may be linked to deviations from typical neurodevelopmental patterns of tNAA. Together, these findings point to tNAA as a promising target for future research on early markers of alcohol-related neural effects as well as and novel treatment targets for youth alcohol use disorders.

## Supporting information

All supplemental figures and tables

## Acknowledgements

This research was supported by National Institute on Alcohol Abuse and Alcoholism grants to LS (NIAAA R21AA030114, NIAAA K23AA025399, K24AA031052). Additionally, AK is supported by NIAAA K01AA031745 grant. We would like to acknowledge the MUSC Computational Brain Imaging Core for neuroimaging data acquisition. TLFB responses were collected and stored via REDCap, which is supported by the National Institutes of Health under Grant Number UL1 TR001450 and hosted by the South Carolina Clinical and Translational Science (SCTR) Institute at the Medical University of South Carolina. We would additionally like to acknowledge the participants and their families for contributing their time and their data to this research.

