## Supplementary material for "Age- and alcohol-related differences in adolescent neurometabolite levels": All supplemental figures and tables

### Supplementary Figures

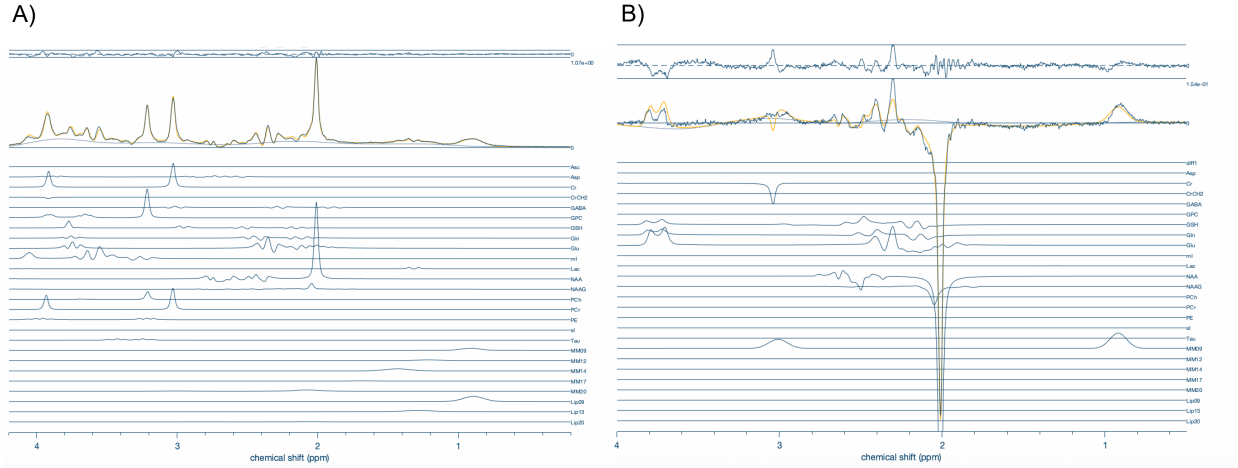

Supplementary Figure 1. Example MR Spectra. A) PRESS sequence for Glx, tNAA, tCho, mI, and tCr. B) MEGAPRESS sequence for GABA+.

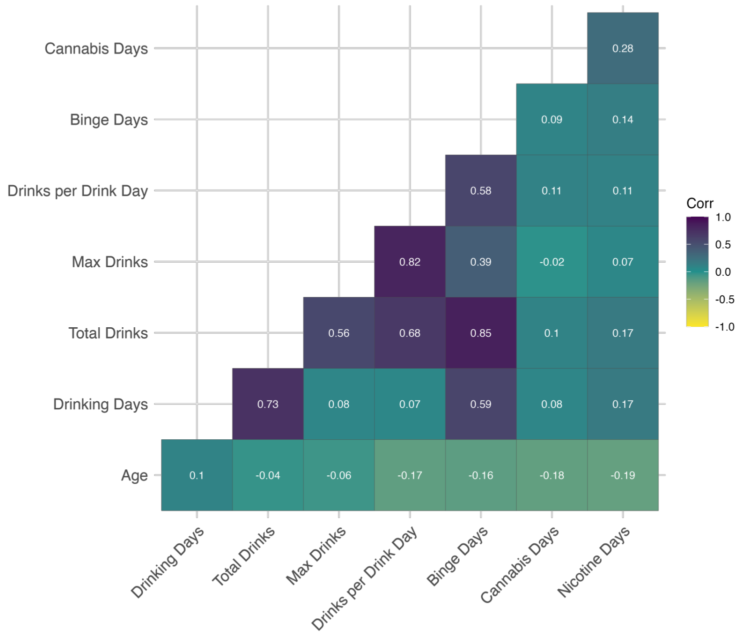

Supplementary Figure 2. Correlations between age and substance use variables.

**Supplementary Tables**

Supplementary Table 1. Full study inclusion and exclusion criteria for each study.

| Study | Cannabidiol (NCT05317546) | N-acetylcysteine (NCT03238300) |
|---|---|---|
| Inclusion criteria | • Ages 16 – 22<br>• Parent or legal guardian informed consent for youth 16 – 17 and youth assent<br>• Informed consent for youth 18 – 22<br>• Meet criteria for AUD in the past year and have at least one current (past 30 days) continued symptom besides craving<br>• Have used alcohol in the past two weeks before screening | • Ages 15 – 19<br>• Parent or legal guardian informed consent for youth 15 – 17 and youth assent<br>• Informed consent for youth 18 – 19<br>• All participants in the alcohol-using group must meet criteria for heavy drinking based on quantity and frequency of drinking (Squeglia et al. 2011; Squeglia et al. 2012)<br>• Non-heavy drinking group must have used alcohol <10 times in their life and has never had a binge drinking episode (4+ females, 5+ males), <5 lifetime experiences with marijuana and none in the past three months; <5 lifetime cigarette use; and no history of other intoxicant use |
| Exclusion criteria | • significant or acutely unstable medical, psychiatric, or substance use problems (e.g., current manic episode, positive psychotic symptoms, severe eating disorder, severe opioid use disorder) that would contraindicate research procedures, interfere with safety, compromise data integrity, or preclude consistent study participation<br>• significant risk of homicide or suicide<br>• currently enrolled in or acutely seeking treatment for AUD or any other SUD<br>• pregnant, trying to become pregnant, or breastfeeding<br>• known allergy or intolerance to CBD<br>• current use of CBD or any supplement containing CBD<br>• history of a serious medical or neurological problem that could affect neural response or brain development | • significant or acutely unstable medical, psychiatric, or substance use problems (e.g., current manic episode, positive psychotic symptoms, severe eating disorder, severe opioid use disorder) that would contraindicate research procedures, interfere with safety, compromise data integrity, or preclude consistent study participation<br>• significant risk of homicide or suicide<br>• currently enrolled in or acutely seeking treatment for AUD or any other SUD<br>• Daily use of tobacco or cannabis<br>• Current DSM-5 diagnosis of moderate or severe substance use disorder (excluding alcohol or cannabis)<br>• Positive urine toxicology screen for narcotics, amphetamines, sedatives, |

| | | |
|---|---|---|
| | • non-correctable visual or hearing problems<br>• MRI contraindications (e.g., braces, claustrophobia, irremovable metal implants or piercings)<br>• Acute drunkenness or consumption of alcohol within 12 hours of visit<br>• Less than or equal to 10 on the Clinical Institute Withdrawal Assessment for Alcohol (CIWA)<br>• Severe Cannabis Use Disorder (CUD)<br>• Concurrent medications with potential drug-drug interactions with CBD, including CYP3A4 and CYP2C19 substrates, inhibitors, and inducers, as well as CYP2C8/9 substrates. | hypnotics, or opiates (unless prescribed)<br>• pregnant, trying to become pregnant, or breastfeeding<br>• Medical conditions or medications that contraindicate N-acetylcysteine (NAC) use<br>• Current use of NAC or any supplement containing NAC (must agree not to use during study participation)<br>• history of a serious medical or neurological problem that could affect neural response or brain development<br>• History of severe asthma not controlled by medication<br>• Less than or equal to10 on the Clinical Institute Withdrawal Assessment for Alcohol (CIWA)<br>• Current use of psychoactive medications that affect cerebral blood flow<br>• Non-correctable visual or hearing impairments<br>• Not fluent in English<br>• MRI contraindications (e.g., braces, claustrophobia, irremovable metal implants or piercings)<br>• Refusal of blood draw<br>• For heavy drinking group, abstinence from alcohol for more than 14 days prior to participation<br>• Use of alcohol within 12 hours prior to scanning (confirmed via breathalyzer) |

Supplementary Table 2. Demographics for each study.

| | Cannabidiol (NCT05317546) | N-acetylcysteine (NCT03238300) |
|---|---|---|
| **Sample size** | 34 | 50 |
| **Age***, range<br>mean (SD) | 16 – 22<br>20.52 (1.48) | 15 – 19<br>18.88 (0.62) |
| **Sex**, n (%)[1]<br>Female<br>Male | <br>25 (74%)<br>9 (26%) | <br>31 (62%)<br>19 (38%) |

| | | |
|---|---|---|
| **Race**, n (%) | | |
| Asian | 1 (3%) | 2 (4%) |
| Black/African American | 4 (12%) | 1 (2%) |
| White | 29 (85%) | 47 (94%) |
| **Ethnicity**, n (%) | | |
| Hispanic/Latino | 2 (6%) | 0 (0%) |
| Not Hispanic/Latino | 32 (94%) | 50 (100%) |
| **Grade*, n (%)** | | |
| 10th grade | 1 (3%) | 1 (2%) |
| 11th grade | 0 (0%) | 8 (16%) |
| 12th grade/GED equivalent | 5 (15%) | 28 (56%) |
| Some college | 21 (62%) | 13 (26%) |
| Bachelor's Degree | 7 (20%) | 0 (0%) |
| **Mental Health Disorders[2], n (%)** | | |
| Attention Deficit Hyperactivity Disorder* | 4 (12%) | 9 (18%) |
| Obsessive-Compulsive Disorder | 0 (0%) | 2 (4%) |
| Major Depressive Episode | 3 (9%) | 1 (2%) |
| Generalized Anxiety Disorder | 6 (18%) | 2 (4%) |
| Social Anxiety Disorder | 3 (9%) | 0 (0%) |
| Panic Disorder | 3 (9%) | 1 (2%) |
| Alcohol Use Disorder | 34 (100%) | 30 (60%) |
| Mild | 19 (56%) | 19 (38%) |
| Moderate | 9 (26%) | 7 (14%) |
| Severe | 6 (18%) | 4 (8%) |
| Cannabis Use Disorder* | 16 (47%) | 16 (32%) |
| Mild | 7 (20%) | 7 (14%) |
| Moderate | 9 (26%) | 3 (6%) |
| Severe | 0 (0%) | 6 (12%) |
| **Alcohol Use[3]** | | |
| Participants with alcohol use in past 60 days n (%) | 34 (100%) | 50 (100%) |
| Number of drinking days, mean (SD) | 13.44 (5.85) | 15.2 (7.21) |
| Total number of standard drinks, mean (SD) | 60.73 (43.89) | 73.76 (48.61) |
| Number of binge drinking days*, mean (SD) | 6.53 (5.07) | 9.48 (7.18) |
| Maximum number of drinks in one day, mean (SD) | 8.57 (4.89) | 8.3 (3.12) |
| Number of standard drinks per drinking day, mean (SD) | 4.62 (2.62) | 4.70 (1.54) |
| Age of participant first full drink | 15.67 (1.94) | 15.47 (1.45) |
| Past 24 hrs self-reported alcohol use | 4 (12%) | 9 (18%) |
| Past 12 hrs self-reported alcohol use | 0 (0%) | 0 (0%) |
| **Other Substance Use[3]** | | |
| Participants with cannabis use in past 60 days | 24 (70%) | 30 (60%) |
| Number of cannabis use days, mean (SD) | 7.71 (12.58) | 12.4 (19.98) |
| Urine tox screen positive for THC | 6 (18%) | 11 (22%) |
| Participants with nicotine[4] use in past 60 days | 16 (47%) | 32 (64%) |
| | 12.82 (20.29) | 20.52 (24.58) |

| | |
|---|---|
| | Number of nicotine use days, mean (SD) |

*Note: Asterisk next to variable name indicates where samples are significantly different (based on t-test or χ² as appropriate). [1]Percents are rounded to the nearest whole number. [2]MINI = Mini International Neuropsychiatric Interview; Current diagnosis was used ensuring that all diagnoses reflected participants conditions at the time of assessment. [3] Based on 60-day Timeline Follow Back (TLFB). Number of substance use days are only quantified using participants that use that substance. [4]Any nicotine use, including tobacco and e-cigarettes.*

Supplementary Table 3. Effective degrees of freedom for GAM models.

| GAM model | edf |
|---|---|
| tNAA ~ s(Age) + GM/BM | 1 |
| tCho ~ s(Age) + GM/BM | 1.56 |
| GABA+ ~ s(Age) + GM/BM | 1.30 |
| mI ~ s(Age) + GM/BM | 1 |
| Glx ~ s(Age) + GM/BM | 1.23 |
| tCr ~ s(Age) + GM/BM | 1 |
| Glx/GABA+ ~ s(Age) + GM/BM | 1 |

Supplementary Table 4. Akaike's Information Criterion (AIC) comparison of linear models and GAM models.

| Metabolite | Model | AIC |
|---|---|---|
| tNAA | Linear model: tNAA ~ Age + GM/BM | 210.27 |
| | GAM model: tNAA ~ s(Age) + GM/BM | 210.27 |
| tCho | Linear model: tCho ~ Age + GM/BM | 38.47 |
| | GAM model: tCho ~ s(Age) + GM/BM | 38.05 |
| GABA+ | Linear model: GABA+ ~ Age + GM/BM | 102.14 |
| | GAM model: GABA+ ~ s(Age) + GM/BM | 102.03 |
| mI | Linear model: mI ~ Age + GM/BM | 278.48 |
| | GAM model: mI ~ s(Age) + GM/BM | 278.48 |
| Glx | Linear model: Glx ~ Age + GM/BM | 332.61 |
| | GAM model: Glx ~ s(Age) + GM/BM | 332.53 |
| tCr | Linear model: tCr ~ Age + GM/BM | 174.96 |
| | GAM model: tCr ~ s(Age) + GM/BM | 174.96 |
| Glx/GABA+ | Linear model: Glx/GABA+ ~ Age + GM/BM | 224.85 |
| | GAM model: Glx/GABA+ ~ s(Age) + GM/BM | 224.85 |

Supplementary Table 5. Statistical significance of covariates tested using likelihood ratio test.

| Predictor | Neurometabolite | Covariate | *p* |
|---|---|---|---|
| Drinking Days | tNAA | Cannabis Days | 0.39 |
| | | Nicotine Days | 0.54 |
| | | Sex | 0.87 |
| | tCho | Cannabis Days | 0.92 |
| | | Nicotine Days | 0.59 |
| | | Sex | 0.21 |
| | GABA+ | Cannabis Days | 0.17 |
| | | Nicotine Days | 0.40 |

| | | | |
|---|---|---|---|
| | | Sex | 0.90 |
| | mI | Cannabis Days | 0.62 |
| | | Nicotine Days | 0.85 |
| | | Sex | 0.46 |
| | Glx | Cannabis Days | 0.62 |
| | | Nicotine Days | 0.28 |
| | | Sex | 0.33 |
| | tCr | Cannabis Days | 0.43 |
| | | Nicotine Days | 0.53 |
| | | Sex | 0.84 |
| | Glx/GABA+ | Cannabis Days | 0.30 |
| | | Nicotine Days | 0.23 |
| | | Sex | 0.46 |
| Total Drinks | tNAA | Cannabis Days | 0.38 |
| | | Nicotine Days | 0.44 |
| | | Sex | 0.69 |
| | tCho | Cannabis Days | 0.90 |
| | | Nicotine Days | 0.60 |
| | | Sex | 0.12 |
| | GABA+ | Cannabis Days | 0.18 |
| | | Nicotine Days | 0.45 |
| | | Sex | 0.79 |
| | mI | Cannabis Days | 0.60 |
| | | Nicotine Days | 0.95 |
| | | Sex | 0.42 |
| | Glx | Cannabis Days | 0.60 |
| | | Nicotine Days | 0.24 |
| | | Sex | 0.34 |
| | tCr | Cannabis Days | 0.44 |
| | | Nicotine Days | 0.52 |
| | | Sex | 0.92 |
| | Glx/GABA+ | Cannabis Days | 0.28 |
| | | Nicotine Days | 0.22 |
| | | Sex | 0.49 |
| Binge Days | tNAA | Cannabis Days | 0.33 |
| | | Nicotine Days | 0.39 |
| | | Sex | 0.83 |
| | tCho | Cannabis Days | 0.95 |
| | | Nicotine Days | 0.57 |
| | | Sex | 0.29 |
| | GABA+ | Cannabis Days | 0.17 |
| | | Nicotine Days | 0.44 |
| | | Sex | 0.92 |
| | mI | Cannabis Days | 0.58 |
| | | Nicotine Days | 0.96 |
| | | Sex | 0.58 |
| | Glx | Cannabis Days | 0.59 |
| | | Nicotine Days | 0.24 |
| | | Sex | 0.27 |

| | | | |
|---|---|---|---|
| | tCr | Cannabis Days | 0.42 |
| | | Nicotine Days | 0.54 |
| | | Sex | 0.79 |
| | Glx/GABA+ | Cannabis Days | 0.30 |
| | | Nicotine Days | 0.23 |
| | | Sex | 0.43 |
| Max Drinks | tNAA | Cannabis Days | 0.32 |
| | | Nicotine Days | 0.28 |
| | | Sex | 0.73 |
| | tCho | Cannabis Days | 0.99 |
| | | Nicotine Days | 0.47 |
| | | Sex | 0.16 |
| | GABA+ | Cannabis Days | 0.17 |
| | | Nicotine Days | 0.41 |
| | | Sex | 0.83 |
| | mI | Cannabis Days | 0.57 |
| | | Nicotine Days | 0.91 |
| | | Sex | 0.76 |
| | Glx | Cannabis Days | 0.57 |
| | | Nicotine Days | 0.21 |
| | | Sex | 0.25 |
| | tCr | Cannabis Days | 0.41 |
| | | Nicotine Days | 0.57 |
| | | Sex | 0.93 |
| | Glx/GABA+ | Cannabis Days | 0.28 |
| | | Nicotine Days | 0.22 |
| | | Sex | 0.49 |
| DPDD | tNAA | Cannabis Days | 0.31 |
| | | Nicotine Days | 0.29 |
| | | Sex | 0.90 |
| | tCho | Cannabis Days | 0.94 |
| | | Nicotine Days | 0.49 |
| | | Sex | 0.14 |
| | GABA+ | Cannabis Days | 0.18 |
| | | Nicotine Days | 0.44 |
| | | Sex | 0.68 |
| | mI | Cannabis Days | 0.53 |
| | | Nicotine Days | 0.92 |
| | | Sex | 0.69 |
| | Glx | Cannabis Days | 0.56 |
| | | Nicotine Days | 0.20 |
| | | Sex | 0.21 |
| | tCr | Cannabis Days | 0.48 |
| | | Nicotine Days | 0.50 |
| | | Sex | 0.77 |
| | Glx/GABA+ | Cannabis Days | 0.30 |
| | | Nicotine Days | 0.22 |
| | | Sex | 0.51 |

*Note: Over the past 60-days, 'Drinking Days' refers to number of days alcohol was consumed. 'Total Drinks' refers to the total number of drinks consumed. 'Binge Days' refers to the number of binge drinking episodes. 'Max Drinks' refers to the maximum number of drinks consumed in one drinking episode. 'DPDD' refers to the average number of drinks per drinking day. Reported p-value refers to the p-value of the likelihood ratio test comparing the base model without that covariate (e.g., tNAA~ drinking_days + age + gm/bm) to a model with that covariate (e.g., tNAA~ drinking_days + age + gm/bm + cannabis_days).*

Supplementary Table 6. Linear model output for associations between metabolites and alcohol variables.

| Metabolite | Predictor | $\beta$ | $\beta$ 95% CI [LL, UL] | t | p |
|---|---|---|---|---|---|
| tCho | *Model: tCho ~ Drinking Days + Age + GM/BM* | | | | |
| | Drinking Days | -0.12 | [-0.33, 0.08] | -1.17 | 0.24 |
| | **Age** | 0.37 | [0.16, 0.57] | 3.56 | 0.00064** |
| | GM/BM | 0.14 | [-0.07, 0.34] | 1.35 | 0.18 |
| | *Model: tCho ~ Total Drinks + Age + GM/BM* | | | | |
| | Total Drinks | -0.16 | [-0.36, 0.05] | -1.53 | 0.13 |
| | **Age** | 0.35 | [0.14, 0.55] | 3.42 | 0.0010** |
| | GM/BM | 0.13 | [-0.07, 0.34] | 1.30 | 0.20 |
| | *Model: tCho ~ Max Drinks + Age + GM/BM* | | | | |
| | Drinks Max | -0.06 | [-0.27, 0.14] | -0.59 | 0.55 |
| | **Age** | 0.35 | [0.14, 0.55] | 3.39 | 0.0011** |
| | GM/BM | 0.15 | [-0.05, 0.36] | 1.50 | 0.14 |
| | *Model: tCho ~ DPDD + Age + GM/BM* | | | | |
| | DPDD | -0.09 | [-0.30, 0.12] | -0.87 | 0.39 |
| | **Age** | 0.34 | [0.13, 0.54] | 3.25 | 0.0017** |
| | GM/BM | 0.15 | [-0.05, 0.35] | 1.47 | 0.15 |
| | *Model: tCho ~ Binge Days + Age + GM/BM* | | | | |
| | Binge Days | -0.20 | [-0.40, 0.01] | -1.91 | 0.06 |
| | **Age** | 0.32 | [0.12, 0.53] | 3.17 | 0.0021** |
| | GM/BM | 0.13 | [-0.08, 0.33] | 1.23 | 0.22 |
| GABA+ | *Model: GABA+ ~ Drinking Days + Age + GM/BM* | | | | |
| | Drinking Days | -0.00 | [-0.20, 0.20] | -0.030 | 0.98 |
| | **Age** | 0.45 | [0.25, 0.65] | 4.51 | 0.000022** |
| | GM/BM | 0.08 | [-0.12, 0.28] | 0.84 | 0.41 |
| | *Model: GABA+ ~ Total Drinks + Age + GM/BM* | | | | |
| | Total Drinks | -0.05 | [-0.25, 0.15] | -0.53 | 0.60 |
| | **Age** | 0.45 | [0.25, 0.65] | 0.78 | 0.000021** |
| | GM/BM | 0.08 | [-0.12, 0.28] | 4.52 | 0.44 |
| | *Model: GABA+ ~ Max Drinks + Age + GM/BM* | | | | |
| | Max Drinks | -0.02 | [-0.22, 0.18] | -0.22 | 0.82 |
| | **Age** | 0.45 | [0.25, 0.65] | 4.51 | 0.000022** |
| | GM/BM | 0.08 | [-0.11, 0.28] | 0.85 | 0.40 |
| | *Model: GABA+ ~ DPDD + Age + GM/BM* | | | | |
| | DPDD | -0.08 | [-0.28, 0.12] | -0.77 | 0.45 |
| | **Age** | 0.44 | [0.24, 0.64] | 4.36 | 0.000038** |
| | GM/BM | 0.08 | [-0.11, 0.28] | 0.84 | 0.41 |
| | *Model: GABA+ ~ Binge Days + Age + GM/BM* | | | | |
| | Binge Days | -0.06 | [-0.26, 0.14] | -0.59 | 0.55 |

| | | | | | |
|---|---|---|---|---|---|
| | **Age** | 0.44 | [0.24, 0.64] | 4.40 | 0.000033** |
| | GM/BM | 0.08 | [-0.12, 0.27] | 0.76 | 0.45 |
| mI | *Model: mI ~ Drinking Days + Age + GM/BM* | | | | |
| | Drinking Days | -0.13 | [-0.33, 0.08] | -1.24 | 0.22 |
| | **Age** | 0.39 | [0.19, 0.60] | 3.83 | 0.00026** |
| | GM/BM | 0.06 | [-0.14, 0.27] | 0.59 | 0.56 |
| | *Model: mI ~ Total Drinks + Age + GM/BM* | | | | |
| | Total Drinks | -0.08 | [-0.28, 0.13] | -0.76 | 0.45 |
| | **Age** | 0.38 | [0.17, 0.58] | 3.67 | 0.00044** |
| | GM/BM | 0.07 | [-0.14, 0.27 | 0.63 | 0.53 |
| | *Model: mI ~ Max Drinks + Age + GM/BM* | | | | |
| | Drinks Max | 0.08 | [-0.12, 0.29] | 0.79 | 0.43 |
| | **Age** | 0.38 | [0.18, 0.59] | 3.74 | 0.00035** |
| | GM/BM | 0.07 | [-0.13, 0.28] | 0.69 | 0.49 |
| | *Model: mI ~ DPDD + Age + GM/BM* | | | | |
| | DPDD | 0.06 | [-0.15, 0.27] | 0.60 | 0.55 |
| | **Age** | 0.39 | [0.18, 0.60] | 3.74 | 0.00035** |
| | GM/BM | 0.08 | [-0.13, 0.28] | 0.73 | 0.46 |
| | *Model: mI ~ Binge Days + Age + GM/BM* | | | | |
| | Binge Days | -0.11 | [-0.32, 0.09] | -1.09 | 0.28 |
| | **Age** | 0.36 | [0.16, 0.57] | 3.50 | 0.00076** |
| | GM/BM | 0.06 | [-0.15, 0.26] | 0.57 | 0.57 |
| Glx | *Model: Glx ~ Drinking Days + Age + GM/BM* | | | | |
| | Drinking Days | 0.10 | [-0.11, 0.30] | 0.96 | 0.34 |
| | **Age** | -0.41 | [-0.62, -0.21] | -4.01 | 0.00013** |
| | GM/BM | 0.01 | [-0.19, 0.22 | 0.14 | 0.89 |
| | *Model: Glx ~ Total Drinks + Age + GM/BM* | | | | |
| | Total Drinks | 0.06 | [-0.14, 0.27] | 0.61 | 0.54 |
| | **Age** | -0.40 | [-0.60, -0.20] | -3.90 | 0.00020** |
| | GM/BM | 0.01 | [-0.19, 0.22] | 0.11 | 0.92 |
| | *Model: Glx ~ Max Drinks + Age + GM/BM* | | | | |
| | Max Drinks | 0.00 | [-0.21, 0.20] | -0.034 | 0.97 |
| | **Age** | -0.40 | [-0.61, -0.20] | -3.91 | 0.00020** |
| | GM/BM | 0.00 | [-0.20, 0.21] | 0.035 | 0.97 |
| | *Model: Glx ~ DPDD + Age + GM/BM* | | | | |
| | DPDD | -0.03 | [-0.23, 0.18] | -0.26 | 0.79 |
| | **Age** | -0.41 | [-0.61, -0.20] | -3.90 | 0.00020** |
| | GM/BM | 0.00 | [-0.20, 0.21] | 0.30 | 0.98 |
| | *Model: Glx ~ Binge Days + Age + GM/BM* | | | | |
| | Binge Days | 0.09 | [-0.12, 0.30] | 0.86 | 0.39 |
| | Age | -0.39 | [-0.59, -0.18] | -3.76 | 0.00033** |
| | GM/BM | 0.02 | [-0.19, 0.22] | 0.15 | 0.88 |
| tCr | *Model: tCr ~ Drinking Days + Age + GM/BM* | | | | |
| | Drinking Days | -0.04 | [-0.26, 0.18] | -0.39 | 0.70 |
| | Age | -0.16 | [-0.38, 0.06] | -1.49 | 0.14 |
| | GM/BM | -0.10 | [-0.32, 0.12] | -0.91 | 0.37 |
| | *Model: tCr ~ Total Drinks + Age + GM/BM* | | | | |
| | Total Drinks | -0.06 | [-0.28, 0.16] | -0.56 | 0.58 |
| | Age | -0.17 | [-0.39, 0.05] | -1.56 | 0.12 |

| | | | | | |
|---|---|---|---|---|---|
| | GM/BM | -0.10 | [-0.32, 0.12] | -0.94 | 0.35 |
| | *Model: tCr ~ Max Drinks + Age + GM/BM* | | | | |
| | Max Drinks | -0.05 | [-0.27, 0.17] | -0.45 | 0.65 |
| | Age | -0.17 | [-0.39, 0.05] | -1.56 | 0.12 |
| | GM/BM | -0.09 | [-0.31, 0.12] | -0.85 | 0.40 |
| | *Model: tCr ~ DPDD + Age + GM/BM* | | | | |
| | DPDD | -0.17 | [-0.38, 0.05] | -1.51 | 0.13 |
| | Age | -0.20 | [-0.41, 0.02] | -1.79 | 0.08 |
| | GM/BM | -0.10 | [-0.31, 0.12] | -0.91 | 0.36 |
| | *Model: tCr ~ Binge Days + Age + GM/BM* | | | | |
| | Binge Days | -0.07 | [-0.29, 0.15] | -0.64 | 0.52 |
| | Age | -0.18 | [-0.40, 0.04] | -1.62 | 0.11 |
| | GM/BM | -0.11 | [-0.33, 0.11] | -0.95 | 0.34 |
| Glx/GABA+ ratio | *Model: Glx/GABA+ ~ Drinking Days + Age + GM/BM* | | | | |
| | Drinking Days | 0.03 | [-0.16, 0.23] | 0.34 | 0.74 |
| | Age | -0.48 | [-0.68, -0.29] | -4.91 | 0.0000048 |
| | GM/BM | -0.04 | [-0.24, 0.15] | -0.45 | 0.65 |
| | *Model: Glx/GABA+ ~ Total Drinks + Age + GM/BM* | | | | |
| | Total Drinks | 0.05 | [-0.15, 0.24] | 0.50 | 0.62 |
| | Age | -0.48 | [-0.67, -0.28] | -4.88 | 0.0000053** |
| | GM/BM | -0.04 | [-0.24, 0.15] | -0.43 | 0.67 |
| | *Model: Glx/GABA+ ~ Max Drinks + Age + GM/BM* | | | | |
| | Max Drinks | 0.03 | [-0.16, 0.23] | 0.33 | 0.74 |
| | Age | -0.48 | [-0.67, -0.28] | -4.87 | 0.0000056** |
| | GM/BM | -0.05 | [-0.24, 0.15] | -0.50 | 0.62 |
| | *Model: Glx/GABA+ ~ DPDD + Age + GM/BM* | | | | |
| | DPDD | 0.04 | [-0.16, 0.24] | 0.42 | 0.68 |
| | Age | -0.47 | [-0.67, -0.28] | -4.76 | 0.0000084** |
| | GM/BM | -0.05 | [-0.24, 0.15] | -0.48 | 0.63 |
| | *Model: tCr ~ Binge Days + Age + GM/BM* | | | | |
| | Binge Days | 0.06 | [-0.14, 0.26] | 0.61 | 0.54 |
| | Age | -0.47 | [-0.67, -0.27] | -4.76 | 0.0000086** |
| | GM/BM | -0.04 | [-0.24, 0.16] | -0.40 | 0.69 |

*Note: 'Drinking Days' refers to number of days spent drinking alcohol over the last 60 days. 'Total Drinks' refers to the total number of drinks consumed over the last 60 days. 'Max Drinks' refers to the maximum number of drinks consumed in one drinking episode. 'DPDD' refers to the number of days per drinking day. 'Binge Days' refers to the number of binge drinking episodes over the last 60 days. $\beta$ indicates standardized coefficients. LL and UL indicate the lower and upper limits of a confidence interval, respectively. Bolded predictor variables represent $p<0.05$. * indicates $p<.05$. ** indicates $p<.01$.*
